# KCC2 Activation Reverses Neurophysiological and Behavioral Deficits in Female Rett Mice

**DOI:** 10.64898/2026.01.13.699303

**Authors:** Muhammad Nauman Arshad, Shu Fun Josephine Ng, Seda Salar, Krithika Abiraman, Toshiya Nishi, Zhong Zhong, Joshua L Smalley, Paul A Davies, Stephen J. Moss

## Abstract

Rett syndrome is an X-linked neurodevelopmental disorder resulting from mutations in the MeCP2 gene, leading to intellectual disability, impaired motor coordination, decreased sociability, and seizures. Central to the underlying pathophysiology are deficits in synaptic inhibition, which are mediated by hyperpolarizing GABA_A_R currents. These events develop postnatally and are dependent upon increased neuronal Cl^-^ extrusion mediated by SLC12A5 (KCC2). Therefore, we tested whether its activation modifies the disease phenotypes evident in female MeCP2^+/-^ mice, using OV350, a direct activator of KCC2. OV350 rapidly induced a sustained reduction in EEG power, accompanied by a decrease in the severity of epileptic discharges. Increased motor coordination, sociability, and spatial memory were also observed. Deficits in KCC2 phosphorylation were also seen in MeCP2^+/-^ mice, consistent with reductions in its activity that were also ameliorated by OV350. Thus, KCC2 activation may be efficacious in limiting the impact of Rett syndrome and other neurodevelopmental disorders.

## 1. Introduction

Rett syndrome is a progressive neurodevelopmental disorder that predominantly affects young girls. The initial clinical symptoms typically emerge between 6 and 18 months of age and include cognitive impairment, repetitive behaviors, respiratory difficulties, and severe seizures.^[1–4]^ Pathological studies have shown that Rett patients exhibit defects in neuronal maturation, characterized by smaller cell bodies, shorter dendrites, fewer synapses, and reduced inhibition, all of which contribute to the occurrence of severe seizures.^[2–4]^ Rett syndrome is linked to various mutations in the Methyl-CpG-binding-protein 2 gene (MeCP2), which plays a critical role in gene regulation and chromatin structure. Due to its DNA-binding capabilities, the absence or reduced functional MeCP2 affects hundreds of target genes, making it challenging to understand how MeCP2 deficiencies cause brain abnormalities in Rett Syndrome. A key gene that plays a critical role in the disease progression and impaired neuronal development in Rett patients is the K+-Cl− co-transporter 2 (KCC2), which is a downstream target gene of MeCP2.^[5]^

KCC2 functions as the primary transporter for chloride extrusion in both developing and mature neurons within the central nervous system.^[6]^ Its activity is critical for the efficacy of fast synaptic inhibition mediated by gamma-aminobutyric acid type A receptors (GABA_A_R), which are ligand-gated ion channels permeable to chloride. The postnatal development of canonical hyperpolarizing GABA_A_R currents reflects a gradual reduction in intraneuronal chloride levels, driven by increased KCC2 expression and its subsequent activity.^7^ The emergence of hyperpolarizing GABA_A_R currents during development is influenced by the phosphorylation status of KCC2, a mechanism that enhances both its membrane trafficking and overall activity.^[7, 8]^ Deficiencies in KCC2 expression and activity disrupt the balance between excitation and inhibition in neural processes, contributing to various brain disorders, including epilepsy, schizophrenia, autism, CDKL5 deficiency disorder, and Rett syndrome.^[9–17]^ Interestingly, KCC2-deficient mice exhibit traits similar to MeCP2 knockout mice, such as impaired motor skills, autism-like behavior, reduced learning capacity, and severe seizures. ^[18–23]^ Both KCC2 and MeCP2 knockout mouse models display an immature neuronal phenotype along with reduced GABAergic inhibition.^[5, 24]^ However, it remains unclear whether enhancing KCC2 activity would alleviate long-term disease phenotypes in a mouse model of Rett Syndrome. To test this possibility, we developed a novel small-molecule activator that specifically potentiates KCC2 activity and effectively terminates pharmaco-resistant seizures. ^[25]^

In this study, we examined the effects of KCC2 activation on neural excitability, inflammation, cognition, sociability, and behavioral phenotypes in MeCP2^+/-^ female mice, a model that closely mirrors the disease characteristics of Rett syndrome. Our results demonstrated that the KCC2 activator, OV350, enhanced cognitive and behavioral outcomes while reducing the severity of epileptic discharges in these mice. These findings suggest that increasing KCC2 activity may effectively restore the disrupted excitatory/inhibitory balance in the brains of those with Rett syndrome, potentially offering symptomatic relief for human patients.

## 2. Results

### 2.1. Rett mice have higher baseline EEG power

Human patients with Rett syndrome exhibit altered resting baseline EEG power compared to typically developing individuals.^[26]^ Specifically, Rett syndrome patients display significantly increased power in the lower frequency delta and theta bands as they age.^[26]^ Considering this, we compared the baseline EEG power between 5-6-month-old WT and MeCP2^+/-^ female mice. To test this, first, we performed EEG surgeries, and one week later, following recovery, we conducted baseline recordings to assess differences in EEG power between the two groups. We analyzed a 20-minute EEG epoch, comparing recordings from both mouse groups. To quantify the differences, we subjected recordings to Fast Fourier transformation (FFT) to convert the EEG signals from the time domain to the frequency domain, thus generating power spectral density plots for frequencies ranging from 0 to 100 Hz (Figure 1A-C). Our findings revealed that the total baseline EEG power of MeCP2^+/^-Veh mice was significantly higher than that of the WT-Veh mice (Figure 1D, WT-Veh: 2.564.75e-007 ± 7.59e-008, n=8; MeCP2^+/-^-Veh: 4.68e-007 ± 1.9e-008, n=9, Welch’s t test, p=0.03). On average, we got a 70% increase in baseline EEG power (Figure 1E, MeCP2^+/-^-Veh: 174.2 ± 7.44% of WT-Veh, n=9, Welch’s t test, p=0.03). To further find that which EEG frequency bands were contributing to an increase in total power, we also assessed the possible differences in the EEG frequency bands (delta; 0–4 Hz, theta; 4–8 Hz, alpha; 8–13 Hz, and beta; 13–30 Hz) and, similar to the human Rett patients, we observed a significant increase in power in the delta and theta frequency bands (Figure 1F, delta: 247.3 ± 22.65% of WT-Veh, Welch’s t test, p=0.02, n=9; theta: 192.1 ± 19.58 of WT-Veh, Welch’s t test, p=0.03, n=9).^[26]^

**Figure 1.**
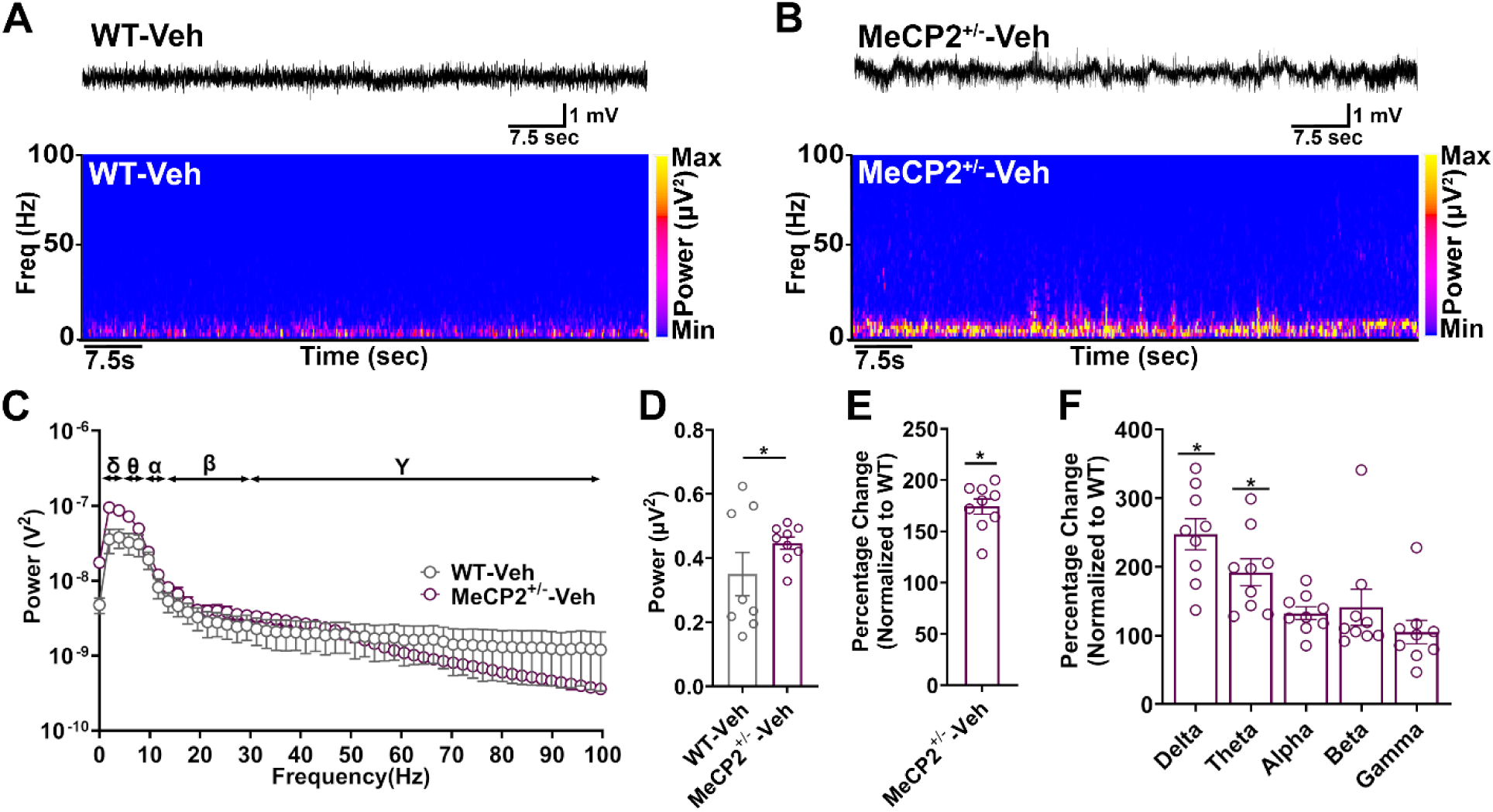
MeCP2 Het females have higher baseline EEG power. **A)** Representative EEG trace and its spectrogram show the power distribution across different frequency bands from WT-Vehicle mice. **B)** Representative EEG trace and its spectrogram show distribution of power across different frequency bands from MeCP2^+/-^-vehicle mice. **C)** EEG recordings from WT-Vehicle and MeCP2^+/-^-vehicle mice were subjected to FFT, and the spectral plot is shown for frequencies between 0 and 100 Hz. **D)** The total EEG power was significantly increased in MeCP2^+/-^ -vehicle mice. **E)** A percentage increase in EEG power was observed in MeCP2^+/-^ - vehicle mice. **F)** The percentage increase in EEG power was observed across the delta and theta frequency bands. Data are presented as mean + SEM. Statistical analysis is performed using Welch’s t-test. ∗p < 0.05.

### 2.2. KCC2 activation reduces total baseline EEG power in Rett mice

Previously, we have shown that KCC2 activation using a novel small-molecule KCC2 activator promotes GABAergic inhibition in the brain.^[25]^ Therefore, next, we compared whether KCC2 activation by an acute dosing with OV350 can alter baseline EEG power in MeCP2^+/-^ mice. To test this, we compared the baseline EEG power, half an hour of the treatment with an acute dose of OV350 (50 mg/kg)/Veh, between both groups of mice (Figure 2A-B). For this, recordings were subjected to an FFT to generate a power spectral density plot for frequencies between 0 and 100 Hz (Figure 2C). The total baseline EEG power of MeCP2^+/-^ -350 mice was significantly reduced compared to the MeCP2^+/-^-Veh mice (Figure 2D, Veh: - 4.415 ± 7.193, n=9; 350: -27.87 ± 7.021, n=10, Welch’s t test, p=0.03). When we compared the power distribution across different frequency bands between OV350 and vehicle-treated mice, we observed that the MeCP2^+/-^ mice treated with OV350 showed a significant reduction in power across the delta and theta frequency bands compared to the vehicle-treated MeCP2^+/-^ mice (Figure 2E, delta: Veh= 23.92 ± 25.81 n=9; 350: -48.35 ± 10.62, n=10, Welch’s t test, p=0.02; theta: Veh: 20.81 ± 10.62, n=9; 350: -26.68 ± 12.62, n=10, Welch’s t test, p=0.01). Next, we compared the effects on the baseline EEG power 18 hours after administration of OV350. For this, recordings were subjected to an FFT to generate a power spectral density plot for frequencies between 0 and 100 Hz (Figure 2F). Interestingly, we observed that KCC2 activation in MeCP2^+/-^ mice treated with OV350 resulted in a lasting impact on baseline EEG power, showing a decrease in total power (Figure 2G, Veh: 23.41 ± 11.44, n=9; 350: -8.519 ± 9.514, n=10, Welch’s t test, p=0.04) as well as in the delta and theta frequency bands, even after 18 hours of treatment (Figure 2H, delta: Veh: 42.62 ± 21.79, n=9; 350: -14.68 ± 14.40, n=10, Welch’s t test, p=0.04; theta: Veh: 16.97 ± 9.3, n=9; 350: -11.76 ± 7.112, n=10, Welch’s t test, p=0.02). This effect is disease-specific, as we didn’t observe any impact of the drug on the total baseline EEG power or across any frequency band when wildtype mice were treated with OV350 (Supplemental Figure S1).

**Figure 2.**
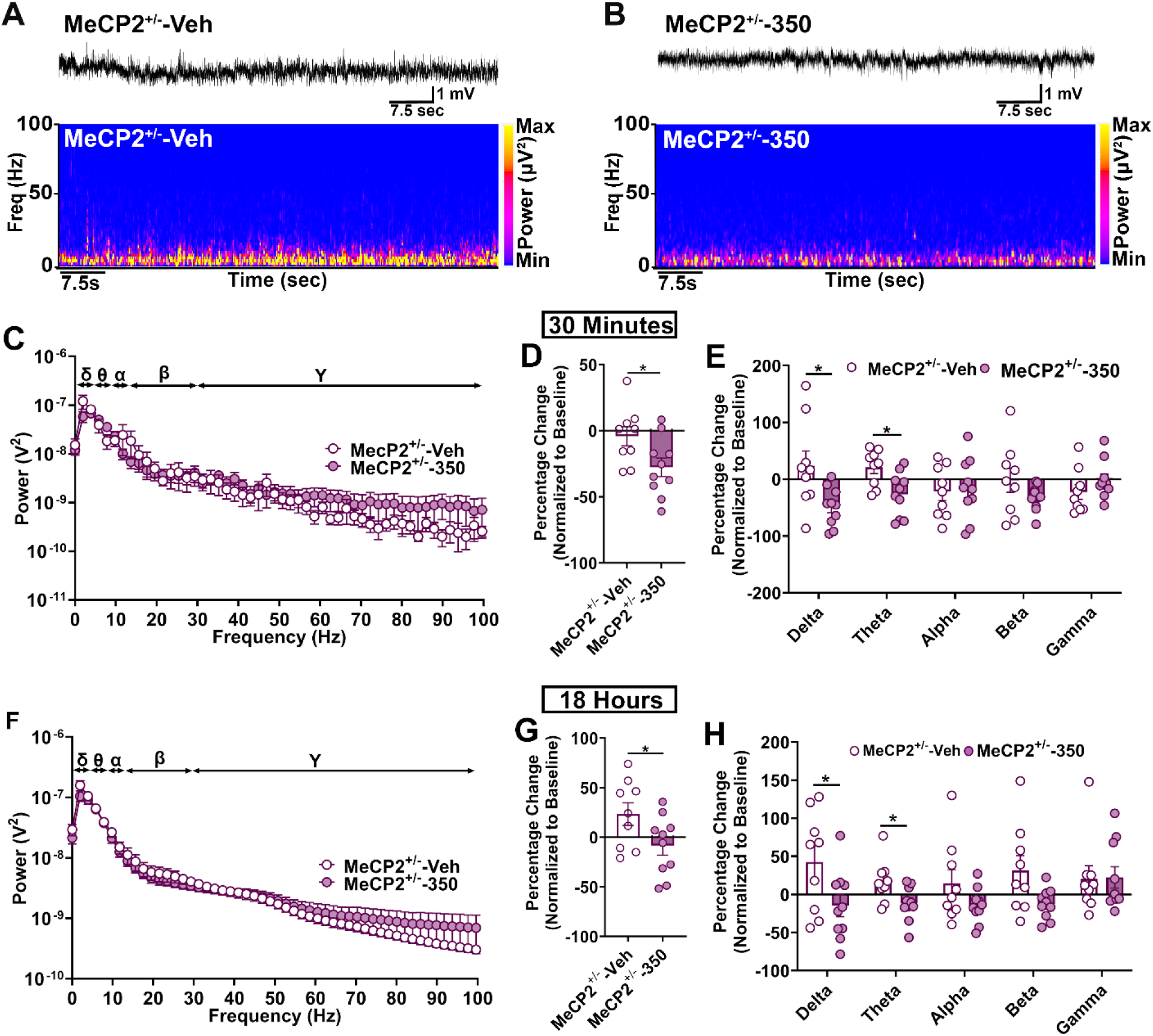
Acute exposure to 50 mg/kg OV350 alters EEG power in MeCP2^+/-^ female mice. **A)** Representative EEG trace and its spectrogram show the power distribution across different frequency bands from MeCP2^+/-^ -vehicle mice. **B)** Representative EEG trace and its spectrogram show power distribution across different frequency bands from MeCP2^+/-^ -350 mice. **C)** EEG recordings from the MeCP2^+/-^ -vehicle and MeCP2^+/-^ -350 mice were subjected to FFT, and the spectral plot is shown for frequencies between 0 and 100 Hz. **D)** The total EEG power was significantly reduced following OV350 treatment in MeCP2^+/-^ mice. **E)** Percentage changes in EEG power were observed across the delta and theta frequency bands following OV350 treatment. We calculated the difference by subtracting the baseline EEG value from the post-treatment EEG value. Next, we divided the difference by the baseline EEG power value and multiplied the result by 100 to express the answer as a percentage change in power. **F)** EEG recordings from 18 hours after the Vehicle/OV350 treatment were subjected to FFT, and spectral plot is shown for frequencies between 0 and 100 Hz. **G)** The total EEG power was significantly reduced 18 hours after the OV350 treatment. **H)** EEG power was significantly altered across the delta and theta frequency bands. Data are presented as mean + SEM. Statistical analysis is performed using Welch’s t-test.∗p < 0.05.

We further examined whether sustained dosing with OV350 impacted the baseline EEG in freely moving mice. We administered a single dose of 50 mg/kg OV350/ Vehicle daily for 5 consecutive days, with recordings continuing for an additional 24 hours after the final dose. To study the effect of repeated doses of OV350 on EEG power, we compared a 20-minute EEG epoch between the two groups of mice on Day 5 (Supplemental Figure S2A). Again, like acute dosing, we observed that OV350 significantly reduced total EEG power in MeCP2^+/-^ mice compared to the vehicle-treated MeCP2^+/-^ mice, and reduced power was also observed across delta and theta frequency bands at 30 minutes (Supplemental Figure S2B-C; Veh: 61.17 ± 20.90, n=9; 350= -25.68 ± -14.26, n=9, Welch’s t test, p=0.03; Supplemental Figure S2D; delta: Veh: 40.89 ± 15.22, n=9; 350: -24.10 ± 16.07, n=9, Welch’s t test, p=0.01, theta; Veh: 28.21 ± 17.01, n=9; 350: -22.98 ± 16.69, n=9, Welch’s t test, p=0.03) and 18 hours after the final dose (Supplemental Figure S2E-F; Veh: 47.63 ± 24.88, n=9; 350: -17.24 ± 11.74, n=9, Welch’s t test, p=0.03; Supplemental Figure S2G; delta: Veh: 31.80 ± 21.22, n=9; 350: -24.79 ± 12.79, n=9, Welch’s t test, p=0.03, theta: Veh: 30.17 ± 17.02, n=9; 350: -5.727 ± 6.812, n=9, Welch’s t test, p=0.04). However, when wildtype mice were treated with OV350 continuously for 5 days, similar to acute exposure, we didn’t observe any impact on total baseline EEG power or across any frequency bands (Supplemental Figure S3). To further confirm that the changes in EEG power were not due to a modified bioavailability of OV350, we compared the distribution of OV350 (50 mg/kg, i.p.) in the plasma and brain of both WT and MeCP2^+/-^ mice. Our results indicated comparable plasma levels of OV350 across genotypes (Supplemental Figure S4A; WT= 23.19 ± 1.976 and MeCP2^+/-^ = 18.47 ± 1.299 (n=4)). Although the blood-brain barrier is compromised in MeCP2^+/-^ mice, we observed no significant difference in the accumulation of OV350 between the genotypes (Supplemental Figure S4B; WT 23.40 ± 3.83 and MeCP2^+/-^ = 24.96 ± 2.645 (n=4)). We also conducted a control experiment to examine the effects of OV350 on locomotion in MeCP2^+/-^ mice using the open field behavioral test. We observed no difference in distance traveled between vehicle-treated MeCP2^+/-^ controls (n=7) and OV350-treated MeCP2^+/-^ mice (n=7), indicating that OV350 does not influence locomotor activity in these mice (Supplemental Figure S4C). Additionally, we measured anxiolytic activity in MeCP2^+/-^ mice and observed no difference in the percentage of time spent in the center between treatment groups, a result consistent with published studies in WT mice (Supplemental Figure S4D).^[25]^

### 2.3. Sustained dosing with OV350 significantly reduces the severity of seizure-like events in Rett mice

Electroencephalographic (EEG) abnormalities, such as epileptic discharges, are also observed in Rett patients at different stages of disease progression.^[27, 28]^ Prevalent types of seizures observed in Rett syndrome patients include complex partial, tonic/atonic, with typical or atypical absence being only infrequently seen.^[29–32]^ This phenotypic co-morbidity is visible in aged MeCP2^+/-^ female mice. ^[33]^

One particularly severe co-morbidity affecting 60–80% of Rett syndrome patients is epileptic seizures, which tend to be poorly controlled by classical anti-convulsant medications.^[29, 34, 35]^ We have shown above that acute and sustained dosing with OV350 reduces baseline EEG power in Rett mice. Therefore, next, we examined whether sustained KCC2 activation using OV350 (50 mg/kg) can suppress seizure-like events in MeCP2^+/-^ mice. Consistent with the literature, we observed epileptic discharges only in the MeCP2^+/-^ mice (Figure 3 A-C). First, we compared the effects of acute dosing on epileptic discharges; we quantified the frequency of epileptic discharges in 30 minutes following 30 minutes of dosing with OV350 on days 1 and 5. We found a comparable number of epileptic discharges in the pretreatment period between the two groups of mice (Figure 3D). On day 1, the OV350 treatment had no effect on the frequency of epileptic discharges (Figure 3E) but significantly reduced their EEG power (Figure 3F, Veh: 11.94 ± 11.74, n=9; 350: -20.83 ± 5.54, n=9, Welch’s t test, p=0.02). We also observed a significant reduction in EEG power across low-frequency bands such as delta and theta (Figure 3G, delta: Veh: 33.49 ± 2.07, n=9; 350: 33.56 ± 10.59, n=9, Welch’s t test, p=0.01; theta: Veh: 16.19 ± 9.165, n=9; 350: -18.20 ± 4.291, n=9, Welch’s t test, p=0.01). Next, we wanted to examine whether sustained dosing of OV350 would reduce the frequency of epileptic discharges in MeCP2^+/-^ female mice. Therefore, on day 5, 30 minutes after the last dose, we compared the frequency and EEG power of events between the two groups of mice and didn’t observe a difference in the frequency of epileptic discharges between the groups (Figure 3H). Interestingly, the EEG power of the events was reduced in OV350-treated MeCP2^+/-^ mice (Figure 3I, Veh: 26.64 ± 9.97, n=9; 350: -17.84 ± 13.65, Mann-Whitney t test, p=0.01). Again, we observed a significant reduction in EEG power across low-frequency bands such as delta and theta in OV350-treated MeCP2^+/-^ mice compared to the vehicle-treated MeCP2^+/-^ mice (Figure 3J, delta, Veh: 19.86 ± 18.48, n=9; 350 -45.17 ± 10.60, n=9, Mann-Whitney t test, p=0.01; theta, Veh: 25.22 ± 9.691, n=9; 350: -18.75 ± 9.303, n=9, Welch’s t test, p=0.01).

**Figure 3.**
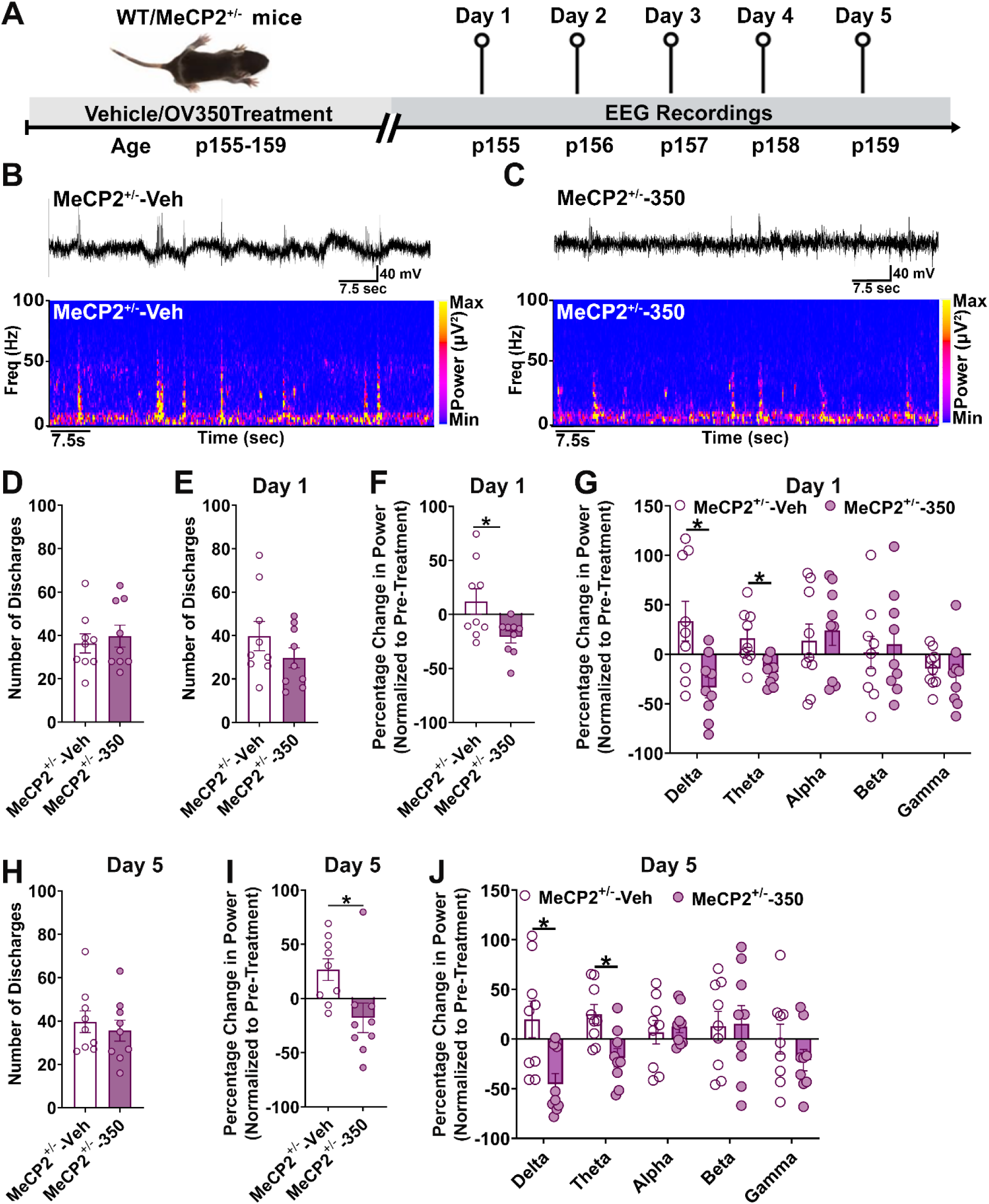
KCC2 activation reduces the intensity of seizure-like events in MeCP2^+/-^ mice. **A)** The line diagram shows the experimental paradigm. **B)** Representative EEG trace and its spectrogram bands from a MeCP2^+/-^ -Vehicle mouse show a series of epileptic discharges. **C)** Representative EEG trace and its spectrogram bands from a MeCP2^+/-^ -350 mouse show a series of epileptic discharges. **D)** Comparison of the frequency of epileptic discharges between the veh and 350-treated MeCP2^+/-^ mice before any treatment. **E)** Comparison of the frequency of epileptic discharges between the veh and 350-treated MeCP2^+/-^ mice 30 minutes after the first treatment. **F)** Percentage change in EEG power of epileptic discharges between the veh and 350-treated MeCP2^+/-^ mice after the first treatment. **G)** Percentage change in EEG power of epileptic discharges across different frequency bands following the first vehicle or OV350 dose. **H)** Comparison of the frequency of epileptic discharges between the veh and 350-treated MeCP2^+/-^ mice 30 minutes after the 5^th^ dose. **I)** Percentage change in EEG power of epileptic discharges between the veh and 350-treated MeCP2^+/-^ mice after the 5^th^ dose. **J)** Percentage change in EEG power of epileptic discharges across different frequency bands following the 5^th^ vehicle or OV350 dose. Data are presented as mean + SEM. Statistical analysis is performed using Welch’s t-test or Mann-Whitney t-test. ∗p < 0.05. The experimental timeline was prepared in BioRender.

### 2.4. OV350 treatment during adulthood alleviates sociability deficits in MeCP2^+/-^ mice

Given that deficits in social interaction are a hallmark feature of autism like behavior and are prevalent in patients with MeCP2 mutations, we next examined sociability using a three-chambered social approach test.^[36]^ To test this, mice received a single dose of OV350 (50 mg/kg)/vehicle daily from p155-162 (Figure 4A-B). During the initial habituation phase, WT and MeCP2^+/-^ mice, irrespective of treatment, explored both chambers equally, demonstrating no initial chamber preference. Consistent with the literature, we found that MeCP2^+/-^ mice demonstrate a profound impairment in the social interaction task, as vehicle-treated MeCP2^+/-^ mice spent significantly less time interacting with the stranger mouse than wild-type mice did (Figure 4C, WT-Veh: 146.4 ± 15.94, n=9; MeCP2^+/-^- Veh: 60.32 ± 12.27, n=8, Welch’s t test, p=0.0007).^[37]^ We also found that the continuous treatment with OV350 (50 mg/kg *i.p*.) significantly improved the time spent with the stranger mouse compared to the vehicle-treated MeCP2^+/-^ mice (Figure 4D, Veh: 60.32 ± 12.27, n=8; 350: 114.2 ± 6.310, n=8, Welch’s t test=0.002). We also found that OV350 treatment had no effect on social interaction in wild-type mice (Supplemental Figure S5A-C). Next, we compared the preference for sociability of all groups of mice. To assess the preference, we compared the time each mouse spent with a dummy or a living mouse. We found that vehicle-treated- WT mice preferred to spend time with a live mouse (Figure 4E, Dummy: 43.96 ± 12.70, n=9; stranger: 146.4 ± 15.94, n=9, paired t test, p=0.0007). Like vehicle-treated WT mice, the OV350-treated WT mice preferred to spend more time with the living mouse (Supplemental Figure S5E, Dummy: 58.06 ± 6.995, n=9; stranger: 119.7 ± 15.96, n=9, paired t test, p=0.001). However, the vehicle-treated MeCP2^+/-^ mice could not differentiate between the live and the dummy mice (Figure 4F). While the KCC2 activation by OV350 in MeCP2^+/-^ mice rescued preference for sociability, as these mice spent more time interacting with the living mouse compared to the dummy mouse (Figure 4G, Dummy: 58.14 ± 12.53, n=8; stranger: 114.2 ± 6.310, n=8, paired t test, p=0.004).

**Figure 4.**
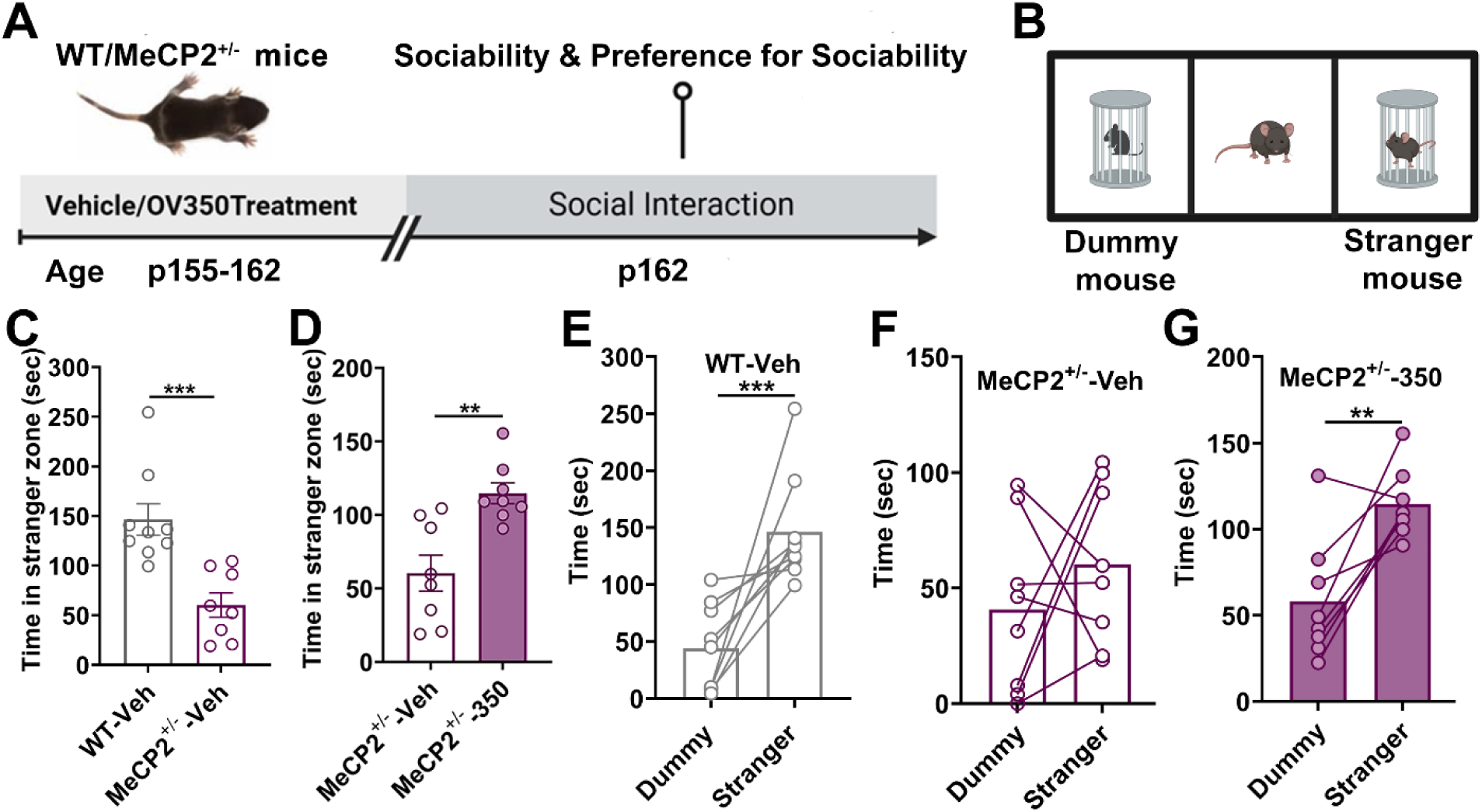
Sustained dosing with OV350 improves sociability in MeCP2^+/-^ mice. **A**) Experimental timeline. The WT and MeCP2^+/-^ mice were administered a single dose of vehicle or OV350 (50 mg/kg *i.p*.) daily from p155-162. We used a three-chamber social interaction assay to assess sociability and preference for social familiarity in the MeCP2^+/-^ mice treated with or without OV350. **B**) Diagram of the sociability assay. Mice were allowed to explore either an unfamiliar mouse or a dummy mouse. **C**) Wild-type mice spent more time interacting with the stranger mouse than vehicle-treated MeCP2^+/-^mice. **D)** OV350-treated MeCP2^+/-^ mice spent more time interacting with the stranger mouse than vehicle-treated mice. **E**) The graph shows that WT-Veh mice preferred interacting with the mouse vs. the dummy mouse. **F**) The graph shows that the vehicle-treated MeCP2^+/-^mice spent equal time with both the dummy and the living mouse. **G)** The graph shows that the 350 treated MeCP2^+/-^mice preferred interacting with the mouse vs. the dummy mouse. Data are presented as mean + SEM. Statistical analysis is performed using Welch’s t-test or paired t-test. ∗p < 0.05, ∗∗p < 0.01, ∗∗∗p < 0.001. The experimental timeline and the cartoon were prepared in BioRender.

### 2.5. Cumulative dosing with OV350 improves learning and memory in Rett mice

To determine whether MeCP2 heterozygotes also have a deficit in learning and memory and whether KCC2 activation can rescue this disease phenotype, wild-type and MeCP2^+/-^ mice were subjected to the novel object recognition task. To test this, we dosed mice daily with OV350 (50 mg/kg)/ vehicle and examined the effects of KCC2 activation on learning and memory in Rett mice (Figure 5A). This task utilizes the differential level of exploration between familiar and unfamiliar (or novel) objects as a behavioral measure of recognition memory, based on the animal’s spontaneous preference for novelty. On day 1, mice are allowed to explore the empty arena for 10 min. On the next day, mice receive a 10-minute training session, during which they are allowed to explore two identical objects placed at symmetric positions from the center of the arena. During the training session, we found that irrespective of the treatment, wild-type and Rett mice performed equally well and spent a similar amount of time on the objects (Figure 5B-E; Supplemental Figure S6A-D). Twenty-four hours post training, mice are reintroduced to the arena and again exposed to two objects, a familiar object and a new object.^[38]^ In this testing phase, we found that compared to the wildtype mice, vehicle-treated MeCP2^+/-^ mice spent less time with the novel object, suggesting deficits in recognition memory (Figure 5F-G, WT-Veh: 60.82 ± 2.48, n=9; MeCP2^+/-^-Veh: 50.13 ± 4.152, n=8, Welch’s t test, p=0.04). Further, we examined whether sustained dosing with OV350 improved recognition in Rett mice and wildtype; to achieve this, we compared the time mice spent interacting with the novel object. We found that OV350-treated MeCP2^+/-^ (Figure 5H) or wildtype (Supplemental Figure S6E-F) didn’t spend more time with the novel object compared to the vehicle-treated controls. Next, we compared whether KCC2 activation improved the preference for the novel object. To test, we compared how much time each mouse spends with the novel object compared to the familiar object, and we found that wild-type mice, irrespective of the treatment, spent more time interacting with the novel object compared to the familiar object (Figure 5I, Familiar: 39.18 ± 2.48, n=9; Novel: 60.82 ± 2.486, n=9, paired t test, p=0.002, Supplemental Figure S6G, WT-Veh: Familiar: 39.18 ± 2.48, n=9; Novel: 60.82 ± 2.486, n=9, paired t test, p=0.002, Supplemental Figure S6H, WT-350: Familiar: 38.98 ± 3.88, n=9; Novel: 61.02 ± 3.88, n=9, paired t test, p=0.02). The vehicle-treated MeCP2^+/-^ couldn’t differentiate between the novel and familiar objects and explored both objects for almost equal time (Figure 5J). Next, we compared the beneficial effects of increasing KCC2 activity by sustained dosing on recognition memory in adult MeCP2^+/-^ mice. We observed that MeCP2^+/-^ mice treated with OV350 preferentially spent more time with the novel object (Figure 5K, Familiar: 40.46 ± 2.83, n=8; Novel: 58.31 ± 2.573, n=8, paired t test, p=0.01). This data suggests that potentiating KCC2 activity by cumulative dosing with OV350 improves recognition memory in MeCP2^+/-^mice.

**Figure 5.**
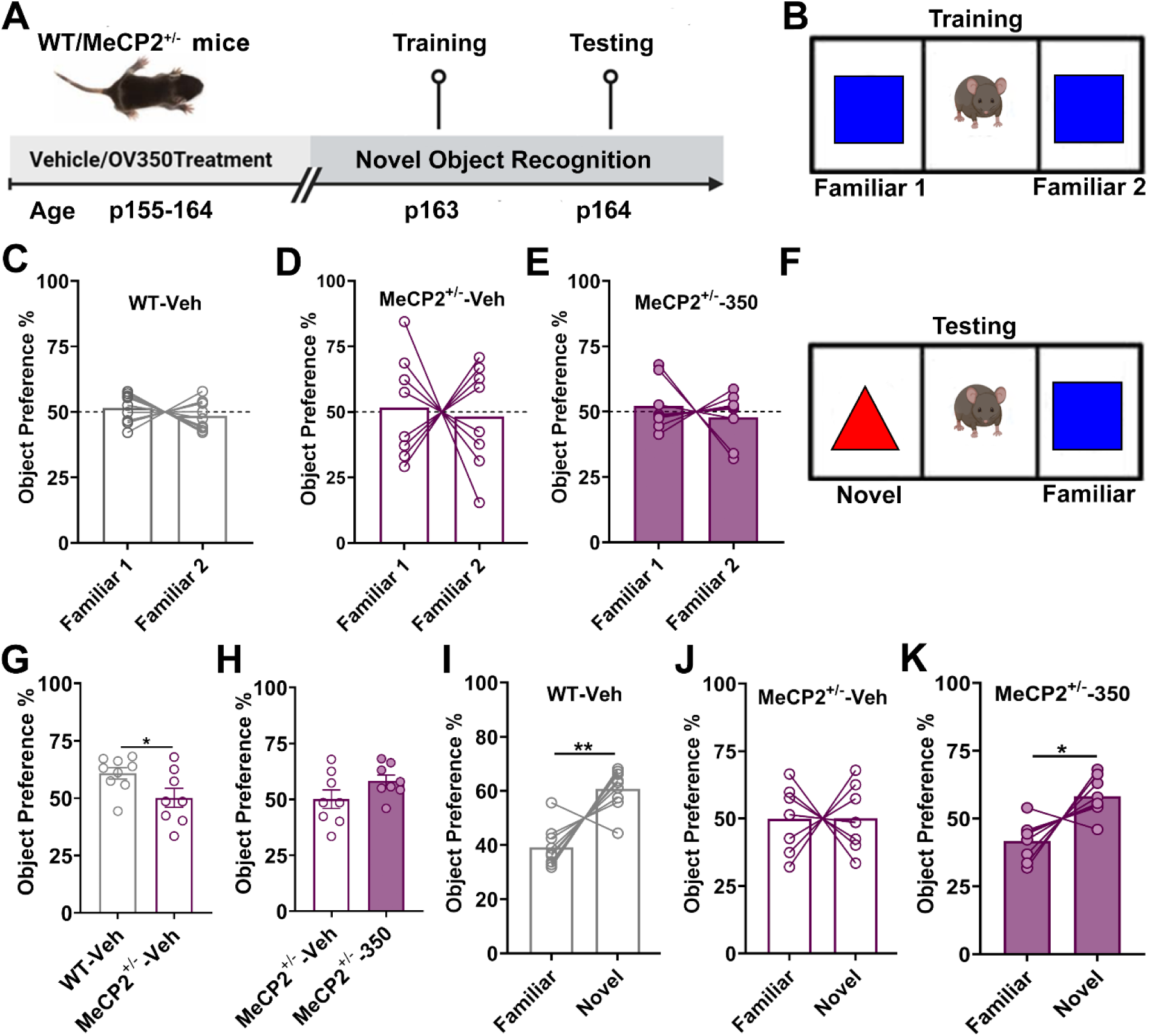
Sustained dosing with 50 mg/kg OV350 improves recognition memory in MeCP2^+/-^ mice. **A)** Experimental timeline. The WT and MeCP2^+/-^ mice were administered a single dose of vehicle or OV350 (50 mg/kg *i.p*.) daily from p155-162. **B)** Representative setup for the training session of the recognition memory task. **C)** The graph shows that WT-Veh spent equal time with both objects. **D)** The graph shows that MeCP2^+/-^-Veh mice spent equal time with both objects. **E)** The graph shows that MeCP2^+/-^-350 mice spent equal time with both objects. **F)** Representative setup for the testing phase of the recognition memory task. **G)** The graph shows that MeCP2^+/-^-Veh mice spent less time with the novel object compared to the WT-Veh mice. **H)** The graph shows that MeCP2^+/-^-Veh and MeCP2^+/-^-350 spent equal time with the novel object. **I)** The graph shows that WT-Veh mice spent more time with the novel object compared to the familiar object. **J)** The graph shows that MeCP2^+/-^-Veh mice couldn’t differentiate between the novel and the familiar object. **K)** The graph shows that MeCP2^+/-^-350 mice spent more time with the novel object compared to the familiar object. Data are presented as mean + SEM. Statistical analysis is performed using Welch’s t-test or paired t-test. ∗p < 0.05, ∗∗p < 0.01. The experimental timeline and the cartoon were prepared in BioRender.

### 2.6. Sustained exposure to OV350 alleviates the severity of hindlimb clasping in MeCP2^+/^ mice

Similar to human patients, MeCP2^+/-^ mice exhibit severe motor deficits, characterized by their inability to fully spread their hindlimbs.^[39]^ Therefore, we examined the effects of continuous dosing with OV350 (50 mg/kg, p155-p164) on hindlimb clasping in these mice. To assess hindlimb clasping scores, we utilized a previously established scale (Figure 6A-B).^[40]^ Consistent with previous studies, vehicle-treated MeCP2^+/-^ mice showed significantly higher hindlimb clasping scores compared to wild-type mice, with 5 out of 8 mice scoring ‘3’ on the scale (Figure 4D, WT-Veh: 0.125 ± 0.125, n=8: MeCP2^+/-^-Veh: 2.25 ± 0.4119, n=8, Mann-Whitney t test, p=0.01). Additionally, we noted that ongoing treatment with OV350 significantly influenced the hindlimb clasping score in MeCP2^+/-^ mice. While the results were not statistically significant, it’s noteworthy that only 1 out of 7 mice scored ‘3’ on the scale (Figure 6E, Veh: 2.25 ± 0.4119, n=8; 350: 1.143 ± 0.404, n=7, Mann-Whitney t test, p=0.07). These findings suggest that while KCC2 activation may lessen the severity of this neurological phenotype, it does not completely prevent it.

**Figure 6.**
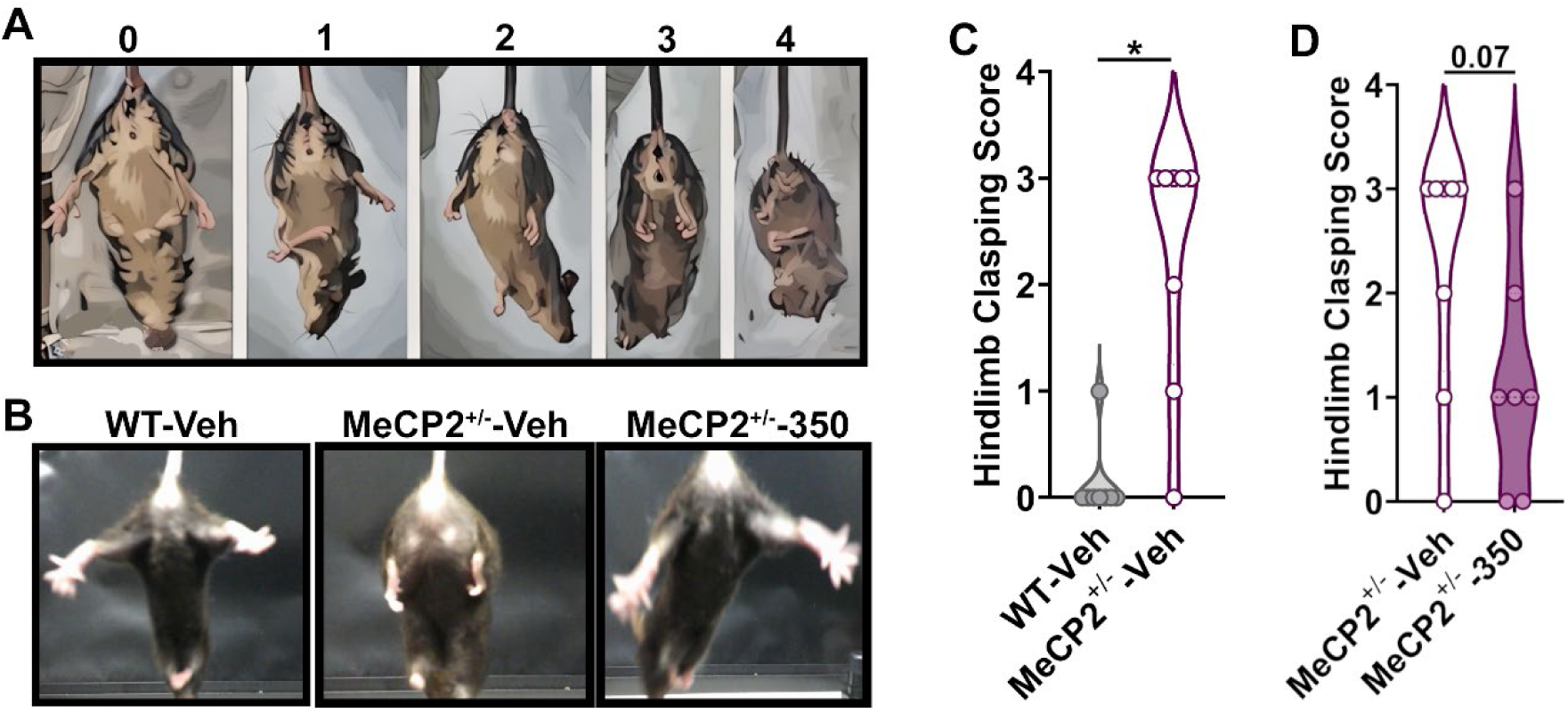
Sustained dosing with OV350 improved hindlimb clasping score in MeCP2^+/-^mice. **A)** The diagram shows different scores of the test based on the severity of the disease phenotype. **B)** Representative images show normal hindlimbs from a WT-Veh mouse, hindlimbs tucked into the abdomen from a MeCP2^+/-^-Veh mouse, and normal hindlimbs from a MeCP2^+/-^-350 mouse. **C**) The graph shows that MeCP2^+/-^-Veh mice show severe hindlimb clasping scores compared to the WT-Veh mice. **D)** The graph shows that OV350-treated MeCP2^+/-^ mice showed considerably improved hindlimb clasping score compared to the vehicle-treated MeCP2^+/-^ mice. Data are presented as mean + SEM. Statistical analysis is performed using the Mann-Whitney t-test. ∗p < 0.05, ∗∗p < 0.01.

### 2.7. KCC2 activation decreases the infiltration of macrophages in the CA1 region of the hippocampus in Rett mice

Neuroinflammation contributes to Rett pathogenesis, as microglia become activated and are subsequently lost with disease progression in animal models of Rett syndrome.^[41–43]^. However, it is unclear whether sustained dosing with OV350 would reduce inflammation in MeCP2^+/-^ mice. Consistent with the literature, we observed a significant reduction in the number of Iba1^+^ cells in MeCP2^+/-^ mice compared to wild-type mice (Figure 7A-B, D, left, WT-Veh: 84.25 ± 3.3, N=12 brain sections; MeCP2^+/-^-Veh: 70.08 ± 2.17, N=12 brain sections, Mann-Whitney t test, p=0.003, right, WT-Veh: 337.0 ± 4.35, n=3; MeCP2^+/-^-Veh: 280.3 ± 11.29, n=3, Mann-Whitney t test, p=0.001).^[42, 43]^ We also found significantly fewer Iba1+&TMEM119+ co-labeled resident microglia in the CA1 area of the hippocampus in MeCP2^+/-^ mice relative to wild-type mice (Figure 7A-B, E, left, WT-Veh: 77.75 ± 3.24, N=12 brain sections; MeCP2^+/-^-Veh: 56.67 ± 2.98, N=12 brain sections, Mann-Whitney t test, p=0.001, right, WT-Veh: 311.0 ± 3.6, n=3 mice; MeCP2^+/-^-Veh: 226.7 ± 12.02, n=3 mice, Mann-Whitney t test, p=0.01). Additionally, there was increased infiltration of Iba1+&TMEM119- co-labeled macrophages in the CA1 area of the hippocampus in MeCP2^+/-^ mice compared to wild-type mice (Figure 7A-B, F, left, WT-Veh: 6.5 ± 0.46, N=12 brain sections; MeCP2^+/-^-Veh: 13.42 ± 1.68, N=12 brain sections, Welch’s t test, p=0.001, right, WT-Veh: 26.00 ± 1.732, n=3; MeCP2^+/-^-Veh: 55.00 ± 7.00, n=3 mice, Welch’s t test, p=0.03). Next, we studied whether KCC2 activation can reverse this disease phenotype in MeCP2^+/-^ mice. Although sustained dosing with OV350 did not change the total number of Iba1^+^ cells in the MeCP2^+/-^ mice (Figure 7B-C, G), we found a higher number of resident glia (Iba1^+^&TMEM119^+^) (Figure 7B-C, H, left, Veh: 70.08 ± 2.17, N=12 brain sections; 350: 75.8 ± 2.7, N=12 brain sections, Mann Whitney t test, p=0.01, right, Veh: 226.7 ± 12.20, n=3 mice; 350: 280.3 ± 12.49, n=3 mice, Welch’s t test, p=0.04) alongside a reduction in the number of macrophages (Iba1^+^&TMEM119^-^) (Figure 7B-C, I, left, Veh: 13.42 ± 1.6, N=12 brain sections; 350: 6.9 ± 0.79, N=12 brain sections, Mann Whitney t test, p=0.01; right, Veh: 55.00 ± 7.00, n=3 mice; 350: 27.67 ± 3.528, n=3 mice, Welch’s t test, p=0.04). This data suggests that repeated administration of OV350 alters the number of active or inflammatory microglia, potentially slowing or preventing Rett pathogenesis.

**Figure 7.**
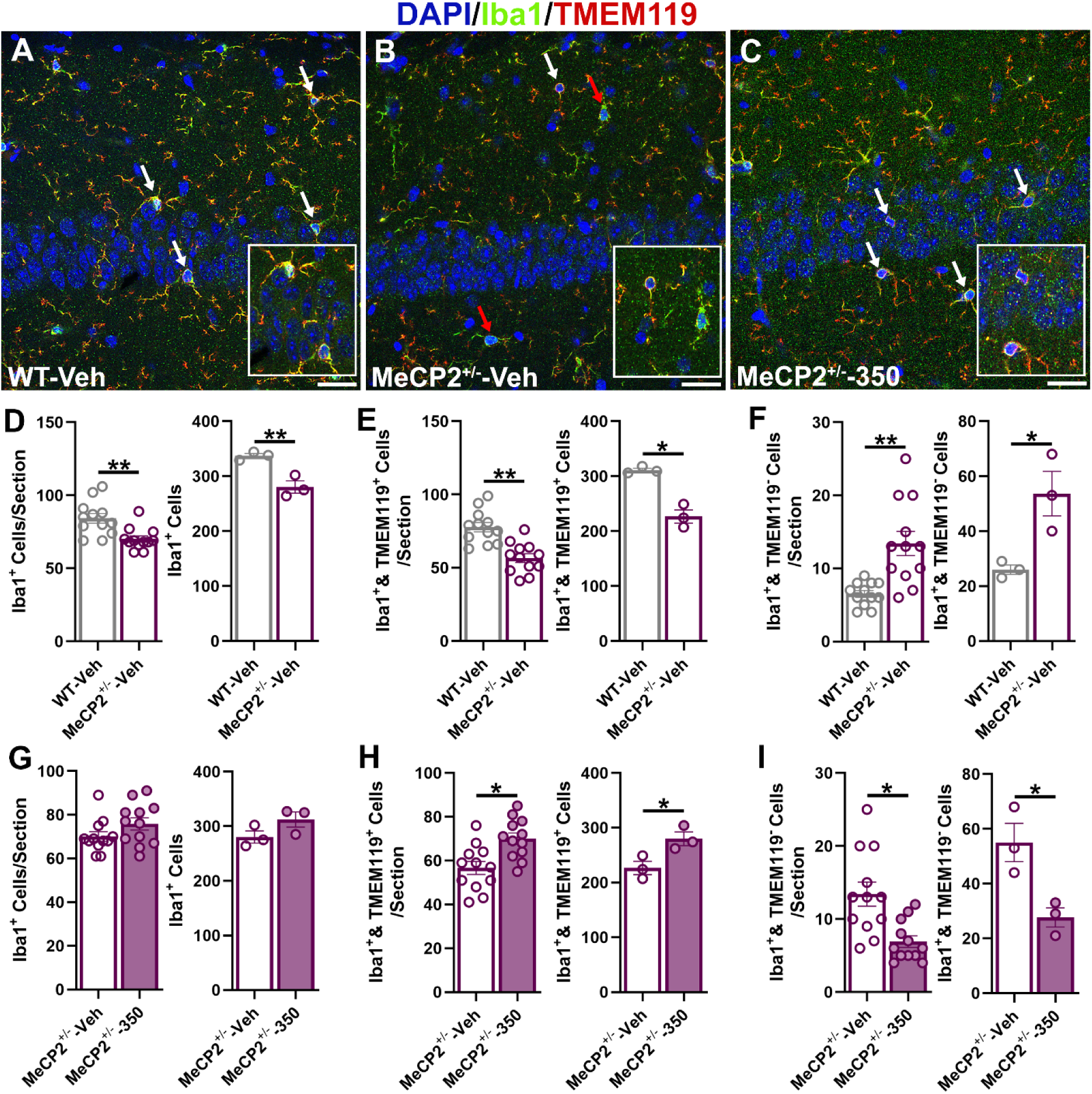
Sustained dosing with OV350 alters the population of macrophages in Rett mice. **A)** Representative image shows Iba1^+^, Iba1^+^ & TMEM119^+^, and Iba1^+^& TMEM119^-^microglia and macrophages, respectively, in a CA1 area of the hippocampus of a WT mouse. White arrows point towards Iba1^+^&TMEM119^+^ co-labeled resident microglia. **B)** Representative image shows Iba1^+^, Iba1^+^ & TMEM119^+^, and Iba1^+^ & TMEM119^-^ microglia and macrophages in a CA1 area of the hippocampus of a vehicle-treated MeCP2^+/-^ mouse. White arrows point towards Iba1^+^ & TMEM119^+^ co-labeled resident microglia. The red arrow points toward Iba1^+^ & TMEM119^-^ macrophage. **C)** Representative image shows Iba1^+^, Iba1^+^ & TMEM119^+^, and Iba1^+^ & TMEM119^-^ microglia and macrophages in a CA1 area of the hippocampus of an OV350-treated MeCP2^+/-^ mouse. White arrows point towards Iba1^+^ & TMEM119^+^ co-labeled resident microglia. **D)** The graph shows significantly fewer Iba1+ cells in the Rett mice. **E)** The graph shows significantly fewer resident microglia in the Rett mice. **F)** The graph shows significantly more macrophages in the Rett mice. **G)** Graph shows no difference in the number of Iba1+ cells in the OV350-treated Rett mice. **H)** The graph shows significantly more resident microglia in the OV350-treated Rett mice. **I)** The graph shows significantly fewer macrophages in the OV350-treated Rett mice. Data are presented as mean + SEM. Statistical analysis is performed using the Welch’s t-test or Mann-Whitney t-test. ∗p < 0.05, ∗∗p < 0.01.

### 2.8. MeCP2 deficiency is associated with aberrant KCC2 phosphorylation and expression

Our results in symptomatic MeCP2^+/-^ mice (5-6 months of age) are consistent with reduced KCC2 activity. Unfortunately, measuring transporter or ion channel function in acute brain slices from older mice using patch clamp recording is fraught with technical limitations that include reduced neuronal viability, increased sensitivity to anoxia, and the complexity of the extracellular matrix, issues that prevent the formation of a high-resistance (Giga-Ohm) seal required for whole-cell patch clamp recordings ^44–46^. Additionally, MeCP2⁺/⁻ female mice, like human Rett syndrome patients, exhibit cellular mosaicism in MeCP2 expression due to the random inactivation of one X chromosome in brain cells. This leads to a heterogeneous population of neurons, some expressing MeCP2 and others lacking it ^47, 48^. Given that MeCP2 regulates KCC2 expression^5^, this mosaicism introduces a significant confound in E_GABA_ measurements to compare KCC2 function between treatments and genotypes. Without knowing whether a recorded neuron is MeCP2-positive or MeCP2-deficient, it becomes exceedingly difficult to interpret the data or draw reliable conclusions.

Given these limitations, we used immunoblotting to measure the phosphorylation of S940 and T1007 within KCC2, which increases and decreases its activity, respectively. These measurements are accepted proxies for KCC2 activity as detailed in published studies from multiple laboratories ^6–8, 16, 49–57^. Compared to WT mice, there was a significant reduction in KCC2 expression in MeCP2^+/-^ mice (Figure 8A-B, WT-Veh: 0.698 ± 0.03, n=4; MeCP2^+/-^- Veh: 0.487 ± 0.03, n=4, Two-way ANOVA, followed by Tukey’s multiple comparison, p=0.01). There was a concomitant increase in the phosphorylation level of KCC2 at the T1007 site (Figure 8A, 8C, WT-Veh: 0.493 ± 0.04, n=4; MeCP2^+/-^- Veh: 0.866 ± 0.03, n=4, Two-way ANOVA, followed by Tukey’s multiple comparison, p=0.01). In contrast, phosphorylation is decreased at S940 (Figure 8A, 8D, WT-Veh: 0.622 ± 0.04, n=4; MeCP2^+/-^-Veh: 0.40 ± 0.03, n=4, Two-way ANOVA, followed by Tukey’s multiple comparison, p=0.01) sites in MeCP2^+/-^ mice. These results demonstrate that reduced MeCP2 expression in Rett mice results in decreased KCC2 expression along with an altered phosphorylation phenotype that displays the hallmarks of an immature phenotype characterized by dysfunctional KCC2 activity, which is strongly correlated with excessive neuronal excitation, seizure-like events, and cognitive and behavioral impairments in adult mice. Next, we assessed whether OV350 treatment rescues KCC2 expression and its phosphoprofile in MeCP2^+/-^ mice. We observed that KCC2 activation in Rett mice doesn’t alter KCC2 expression (Figure 8A-B). We also found that OV350 treatment reduces phosphorylation at T1007 (Figure 8A, 8C, Veh: 0.866 ± 0.03, n=4; 350: 0.61 ± 0.05, n=4, Two-way ANOVA, followed by Tukey’s multiple comparison, p=0.01) while increasing the phosphorylation at the S940 site in MeCP2^+/-^ mice (Figure 8A, 8D, Veh: 0.407 ± 0.03, n=4; 350: 0.58 ± 0.03, n=4, Two-way ANOVA, followed by Tukey’s multiple comparison, p=0.03). These findings suggest that OV350 potentiates KCC2 activity in MeCP2^+/-^ mice without modifying its expression but alters the phosphorylation of the key regulatory residues S940 and T1007.

**Figure 8.**
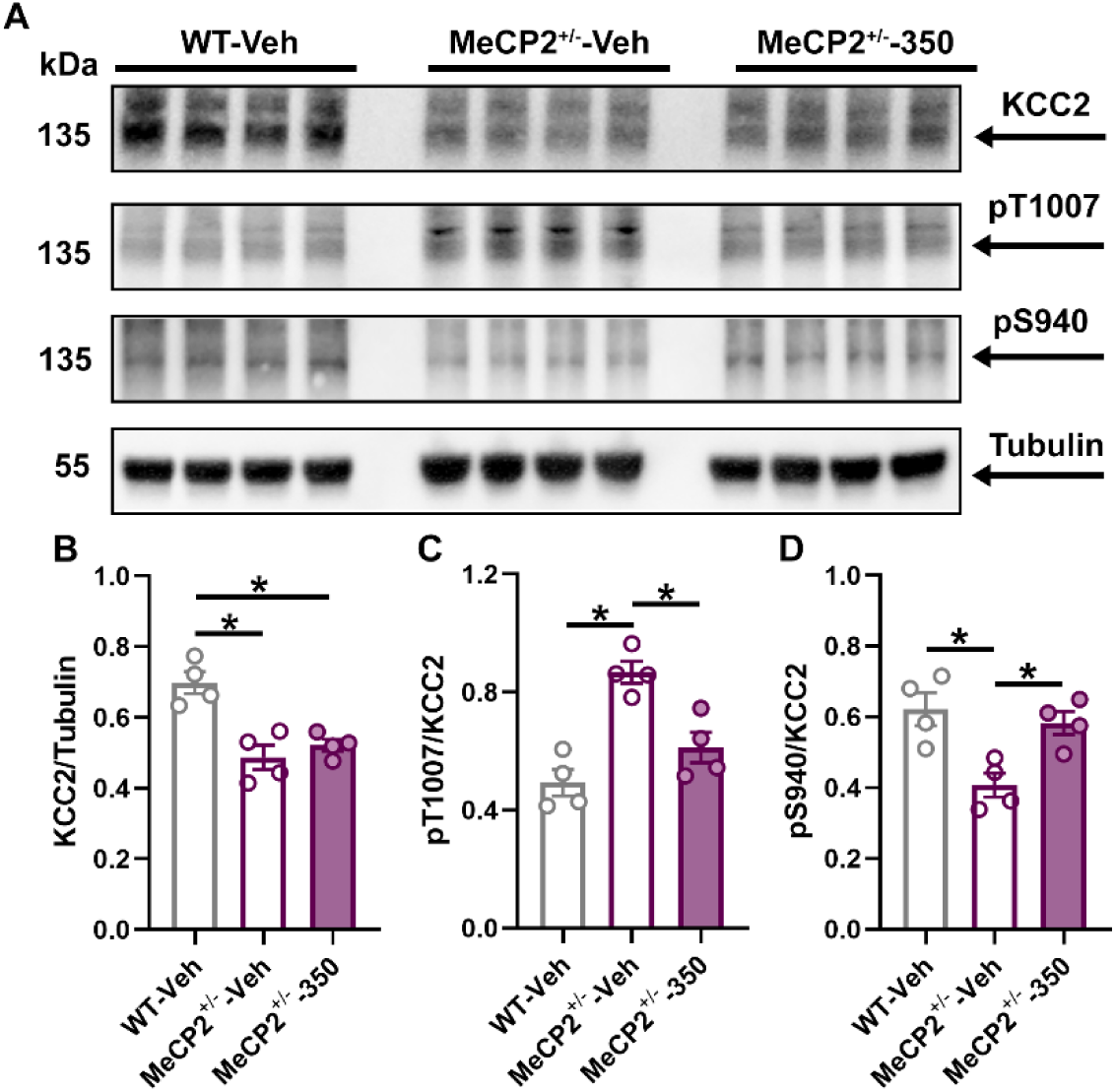
Sustained dosing with OV350 alters the phosphoprofile of KCC2 in MeCP2^+/-^-mice. **A)** Forebrain lysates prepared from the WT-Veh, MeCP2^+/-^-Veh, and MeCP2^+/-^-350 mice were immunoblotted with beta-actin, KCC2, and phospho-specific antibodies against S940(pS940), and T1007(pT1007). **B)** The graph shows a reduction in KCC2 expression in MeCP2^+/-^-Veh, and MeCP2^+/-^-350 mice compared to the WT-Veh mice. **C)** KCC2 phosphorylation at T1007 is increased in MeCP2^+/-^-Veh compared to the WT-Veh mice. Sustained dosing with OV350 reduced phosphorylation at the T1007 site in MeCP2^+/-^-mice compared to the vehicle-treated MeCP2^+/-^ mice. **D)** S940 phosphorylation of KCC2 is reduced in MeCP2^+/-^-Veh compared to the WT-Veh mice. Sustained dosing with OV350 increased phosphorylation at the S940 site in MeCP2^+/-^-mice compared to the Veh-treated MeCP2^+/-^ mice. Data are presented as mean + SEM. Statistical analysis is performed using the Two-Way ANOVA, followed by Tukey’s Multiple Comparison. ∗p < 0.05.

## 3. Discussion

Pharmacological activation of KCC2 represents a promising strategy for enhancing GABAergic inhibition in the brain, offering a therapeutic approach for alleviating disease phenotypes associated with Rett Syndrome. In our study, we found that direct activation of KCC2 normalizes baseline EEG power, reduces the severity of epileptic discharges, and diminishes disease-associated neuroinflammation in MeCP2^+/-^ female mice. Furthermore, our findings indicate that sustained dosing with a KCC2 activator improves sociability, cognitive function, and behavioral deficits in Rett mice.

KCC2 functions by extruding intracellular chloride anions and is essential for the ontogenetic switch of GABAA-mediated responses from depolarizing to hyperpolarizing.^[6, 7]^ The expression and function of KCC2 play a crucial role in regulating neuronal excitability, and disruptions in KCC2 have been associated with an early onset of epilepsy and cognitive deficits.^[20]^ Our findings indicate a significant decrease in KCC2 expression in cortico-hippocampal samples from MeCP2^+/-^ mice, which aligns with current literature on altered KCC2 expression in human Rett patients and animal models of Rett syndrome.^[58,59]^ This reduction in expression may occur because KCC2 is a critical downstream gene target of MeCP2, suggesting that rescuing KCC2 expression using insulin-like growth factor 1 in MeCP2-deficient neurons could restore GABAergic function.^[5]^

It is well established that phosphorylation of KCC2 at the serine 940 site (pS940) is critical for stabilizing KCC2 on the neuronal membrane surface, enabling the extrusion of Cl-and maintaining KCC2 activity.^[52]^ We observed a significant reduction in phosphorylation at the S940 site on KCC2 in the total protein lysates from MeCP2^+/-^ mice. Additionally, we identified a significant increase in phosphorylation at the threonine 1007 site on KCC2 (pKCC2-T1007), which inhibits its transport function.^[54]^ This alteration may result from the inactivation of the mTOR signaling pathway, which is essential for the phosphorylation of KCC2.^[60]^ Our findings of altered phosphorylation suggest a potential delay in this critical developmental switch, which may contribute to cognitive and behavioral deficits, hyperexcitability, and infantile seizures in both human patients and animal models of Rett syndrome. Our findings suggest that OV350 treatment enhances GABAergic inhibition. This is accompanied by an increase in KCC2 phosphorylation at the S940 site and a corresponding decrease in phosphorylation at the T1007 site, which is consistent with its activation and enhanced accumulation on the plasma membrane. These effects may reflect reduced NMDA receptor activity, leading to PP1-dependent dephosphorylation of S940.^[49]^ Additionally, these changes might coincide with a decrease in WNK-dependent phosphorylation of T1007, a process known to diminish KCC2 activation and cause its internalization.^[57]^ These findings suggest that OV350 may help maintain the balance between excitation and inhibition by enhancing KCC2 activity and stability in MeCP2-deficient female mice.

Neuronal and circuit hyperexcitability has been observed in EEG recordings from patients with Rett syndrome, as well as in EEG and brain slice recordings from Rett mice, and in multielectrode recordings of neurons derived from Rett patient organoids.^[26,33,61]^ To investigate this phenomenon, we compared baseline EEG power in MeCP2^+/-^ mice with that of wild-type mice. Our analysis revealed a significant increase in baseline EEG power among the Rett mice. This finding aligns with EEG studies conducted on human patients, where we noted elevated EEG power across the delta and theta frequency bands.^[26]^ These results suggest that quantitative EEG measurements may serve as reliable biomarkers for the accurate diagnosis of Rett syndrome and for the effective development of new treatments. Furthermore, we discovered that administering a KCC2 activator during postnatal development was sufficient to normalize EEG power levels in adult MeCP2^+/-^ mice. Currently, there is a lack of validated biomarkers to assess brain function and clinical severity in individuals with Rett syndrome, complicating the objective evaluation of emerging treatments. Therefore, we advocate for the use of quantitative EEG parameters as objective measures of brain function and disease severity in future clinical trials for Rett syndrome.

Both the rodent model of Rett syndrome and human patients show drug-resistant seizures.^[33,30, 62]^ Our research indicates that MeCP2^+/-^ mice exhibit a rhythmic pattern of epileptic discharges that are absent in WT mice, potentially due to their elevated basal EEG power and deficits in KCC2 activity. Notably, we discovered that repeated administration of OV350 reduces the severity of the epileptic discharges in the MeCP2^+/-^ mice. We propose that restoring the balance of excitation and inhibition by enhancing GABAergic inhibition through rescuing KCC2 activity may serve as an effective therapeutic strategy to prevent drug-resistant seizures in Rett patients.

Major autistic-like phenotypes, including social and communication deficits, are defining characteristics of Rett syndrome.^[63]^ Consistent with existing research, our findings demonstrate that MeCP2^+/-^ mice exhibited significantly poorer performance in social interaction tasks compared to their wild-type counterparts. Notably, we observed an improvement in sociability deficits when we enhanced KCC2 activity through sustained dosing with OV350. This approach may facilitate the transition from excitation to inhibition within neural circuits, thereby achieving the necessary GABA-mediated inhibition to improve this behavioral phenotype in MeCP2^+/-^ mice. These results underscore the significance of KCC2 as a potential therapeutic target for alleviating sociability deficits associated with Rett syndrome.

Mice with reduced or absent MeCP2 expression display several significant characteristics associated with neurological disorders, including impairments in hippocampal-dependent memory and deficits in motor coordination.^[64,65]^ While both young adult and middle-aged female Rett mice demonstrate learning and memory deficits, they typically do not show motor impairments at an early age. Consistent with these observations, our studies revealed learning and memory impairments in MeCP2^+/-^ female mice. Notably, we found that enhancing KCC2 activity in these MeCP2^+/-^ mice resulted in improvements in hippocampal-dependent learning and memory, leading to better performance on the novel object recognition task. This is in line with our previous research, which indicated that the constitutive activation of KCC2 through the development of a transgenic mouse line enhanced learning and memory.^[7]^ Furthermore, recent studies involving Rett mouse models have also found that increasing KCC2 expression can reduce motor deficits. Therefore, our findings reinforce the idea that targeting KCC2 activity may represent the most effective strategy for mitigating cognitive deficits in Rett mice.

In line with existing literature, we observed a reduced number of Iba1⁺ cells and an increase in the population of infiltrating Iba1⁺/TMEM119⁻ macrophages in MeCP2-deficient mice.^[42]^ Following KCC2 activation through OV350 treatment, there was a noted decrease in these infiltrating Iba1⁺/TMEM119⁻ macrophages. The exact mechanism by which KCC2 activation alleviates neuroinflammation remains unclear. Previous studies have indicated that MeCP2 plays a role in regulating the expression of proinflammatory genes, including IL-1β, IL-6, TNF-α, and CX3CR1.^[42, 66, 67]^ Mutations in MeCP2 are associated with neuroinflammation that may worsen the pathology and severity of related disorders, including seizure-like episodes, cognitive deficits, and autism-like behaviors. Notably, MeCP2 deficiency has been shown to enhance the expression of TNFα and other cytokines through increased NF-κB signaling.^[66]^ It is known that TNFα, in turn, downregulates the activity and expression of the neuronal KCC2, and our findings in this study, along with those of others, have reported decreased KCC2 expression in MeCP2-deficient mice.^[68–70]^ This interaction represents an important mechanism through which neuroinflammation can lead to neuronal hyperexcitability, contributing to disorders such as epilepsy and autism spectrum disorder—phenotypes that are also characteristic of Rett syndrome. Therefore, the enhancement of KCC2 activity via OV350 may possess anti-inflammatory properties that could positively influence seizure activity and autism-like behaviors. Additionally, it remains to be clarified whether OV350 affects the activation state of microglia, an area warranting further investigation in future studies.

Several important limitations should be considered before generalizing these findings to patient populations. First, our study focused exclusively on a single dose of OV350 (50 mg/kg), which has previously been shown to reverse diazepam-resistant seizures in adult mice and effectively activate KCC2 for at least 8 hours following intraperitoneal injection. We plan to use varying dosages to achieve disease-modifying effects in Rett syndrome mice. Second, our treatment was administered over several days, and it remains to be seen whether a more condensed treatment schedule would yield similar effectiveness. Third, we used only heterozygous MeCP2 female mice in this study, and it is uncertain whether OV350 treatment would have disease-modifying effects in male MeCP2-deficient mice. Conducting this research will be challenging, as MeCP2-deficient male mice typically do not survive beyond 6 to 8 weeks of age. Nevertheless, future experiments are essential to investigate whether OV350 can reduce disease-related phenotypes in male Rett syndrome mice. Fourth, in our study, MeCP2-deficient female mice remained on OV350 throughout all behavioral and EEG experiments. It is important to determine whether phenotypic improvements persist after treatment cessation or if ongoing dosing is necessary to maintain a rescue of disease symptoms. Lastly, to validate OV350 as a KCC2 activator, it is important to combine OV350 with a KCC2 antagonist and assess whether OV350 directly activates KCC2 and demonstrates a disease-modifying effect in Rett syndrome.

In conclusion, we observed that the reduced MeCP2 expression results in aberrant phosphorylation of KCC2. We also found that MeCP2^+/-^ female mice have higher baseline EEG power, exhibit epileptic discharges, neuroinflammation, and show poor sociability and deficits in learning and memory compared to WT mice. These deficits in adult mice are negated by potentiating KCC2 function. This study demonstrates that enhancing KCC2 function using a novel small-molecule activator is a potential therapy for Rett syndrome and other developmental and epileptic encephalopathies and supports further study.

## Materials and Methods

### Study Design

This study tested whether the intervention with the KCC2 activator, OV350, would correct phenotypes seen in adult MeCP2^+/-^ mice. We selected three phenotypes to test that correspond to clinical phenotypes observed in Rett patients, namely, altered electroencephalographic activity, seizure-like events, recognition memory impairments, and behavioral deficiencies. All experiments included age-matched female mice, with litters randomized to treatment with vehicle or OV350. Outliers were tested for, but none were found; hence, no data were excluded from the study.

### Animals

The TUFTS University Institutional Animal Care and Use Committee approved all animal use. The animals were housed in temperature-controlled rooms on a 12-hour day/night cycle. We purchased the MeCP2^+/-^ mice from The Jackson Laboratory (Strain# 003890), and littermate wildtype females were used for our experiments.

### Drug preparation

OV350 was formulated with 6.25% DMSO and 93.75% (v/v) of 50% (w/v) Captisol, and the Vehicle is 6.25% DMSO and 93.75% (v/v) of 50% (w/v) Captisol. Mice were injected with 50 mg/kg OV350 (i.p., intraperitoneal), a dose which has previously been shown to reach a brain concentration of 676 nM within 4 hours and is maintained for 8 hours.^[25]^

### EEG surgeries and recording

Adult mice were subjected to EEG surgeries as previously described.^[71]^ After the surgery, the mice recovered for seven days in their home cages before being used for experimentation. On the day of recording, the mice were connected to the pre-amps for recording. First, a ∼ 20-hour-long baseline recording was obtained; subsequently, the mice received a single dose of 50 mg/kg OV350 daily for five consecutive days. Following that, the experiment concluded, and the mice were disconnected and returned to their home cage.

### EEG analysis

To assess the potential impact of OV350 treatment on baseline EEG power, a 20-minute silent period was analyzed, during which no muscular movement was detected based on the EMG channel. This represented the pretreatment measurement. Mice received a single dose of 50 mg/kg OV350 or vehicle control for 5 consecutive days. The effects of OV350 on baseline EEG power were assessed 30 minutes and 18 hours after injections on Days 1 and 5 during a 20-minute-long posttreatment period. The percentage change in power was statistically compared between the treatment groups. To compare the EEG signals, they were transformed into the frequency domain using the fast Fourier transform (FFT), producing power spectral plots using the LabChart software. This analysis used an 8 K FFT size, Hann (cosine-bell) window, and 87.5% window overlap. The EEG frequency analysis involved binning the signals into different frequency bands, including delta (0–4 Hz), theta (4–8 Hz), alpha (8–13 Hz), beta (13–30 Hz), and gamma (30–100 Hz). The contribution of each band to the total power was then calculated and compared across treatment groups. A seizure-like event was defined as EEG activity 2.5 times the standard deviation of the preceding 1 min of activity.^[72]^

### Brain Exposure

Mice were administered OV350 intraperitoneally (i.p.) in 6.25% DMSO and 93.75% (v/v) of 50% (w/v) Captisol, with a maximum injection volume of 300 μL. Brain and plasma samples were collected one hour post-treatment from both WT and MeCP2^+/-^ mice, and were quickly frozen on dry ice. Subsequently, plasma and tissue samples were extracted using acetonitrile, and the solvent phases were removed in preparation for LC-MS/MS analysis (Oakland Analytics, Inc., Berkeley, CA 94710; www.oakland-analytics.com).

### Behavior

For all behavioral tests, WT and MeCP2^+/-^ female mice were given injections of either a vehicle or OV350. Littermates were consistently tested at the same time. After each experiment, the equipment was sanitized after each mouse using 70% ethanol, followed by Clidox. Only female mice were utilized for all experiments.

### Open field

Individual mice were positioned in the center of a 40 cm × 40 cm arena, and their movements were monitored using a photobeam system (Motor Monitor, Kinder Scientific). MeCP2^+/-^ mice received either a vehicle or OV350 (intraperitoneal injection at 50 mg/kg). One hour after the injections, the mice were placed in the center of the arena, and the total distance traveled in the open field was compared to that of the vehicle-treated control group. The total distance traveled over a duration of 10 minutes and the percentage of time spent in the center were measured.

### 3-Chamber Social Interaction Assay

Mice were tested in a 3-chamber setup, with each chamber measuring 40 cm × 40 cm. The test mice were allowed to explore the arena for 5 minutes for habituation. After this, an unfamiliar age-matched female mouse was placed in one of the cages, and a dummy mouse was placed in the second cage. The test mice were then allowed to explore the arena for 10 minutes. The time spent in the chamber with the familiar versus the dummy mouse was also calculated. An overhead camera and Ethovision software were used to detect time spent in each region of the arena.

### Novel Object Recognition Test

Age-matched WT and MeCP2-deficient female mice were used for the assay. Mice were assessed using a 3-chamber setup, with each chamber measuring 40 cm × 40 cm. For habituation, mice were allowed to freely explore the experimental arena for 10 minutes, during which no objects were present. On the following day, the mice were introduced into the arena, where they encountered two identical objects and were allowed to explore for 10 minutes (training phase). Twenty-four hours after this training session, one of the objects was substituted with a novel object that differed in shape and color from the familiar ones. The mice then explored the arena for another 10 minutes. Exploration was defined as sniffing or touching the objects with the nose and/or forepaws. An overhead camera, along with the Ethovision software, was used to detect time spent interacting with objects.

### Hindlimb clasping

Hindlimb clasping is a marker of disease progression and neurodegeneration, including Rett Syndrome.^[40, 73]^ To perform this behavioral task, we grasped the tail near its base and lifted the mouse clear of all surrounding objects. Hindlimb position was observed for ∼10 seconds. 1) If the hindlimbs were spread outward, away from the abdomen, it was assigned a score of 0. 2) If one hindlimb was retracted toward the abdomen for more than 50% of the time suspended, it received a score of 1. If both hindlimbs were partially retracted toward the abdomen for more than 50% of the time suspended, it received a score of 2. 3) If both hindlimbs of mice were entirely retracted and touching the abdomen for more than 50% of the time suspended, it received a score of 3. If all four limbs were retracted toward the abdomen for more than 50% of the time suspended, it received a score of 4. Mice were returned to their home cage after recording the hindlimb clasping score.

### Immunohistochemistry

Mice were perfused with 4% paraformaldehyde in 0.1 M sodium phosphate buffer (pH 7.4), and brains were removed from the skull, postfixed for 15 h at 4 °C, and equilibrated in ascending sucrose solutions (10%, 20%, and 30% sucrose in 0.1 M sodium phosphate buffer, pH 7.4) as described previously.^[74]^ Forty μm cryostat sections in the horizontal plane were collected in 0.1 M sodium phosphate buffer saline (PBS). Immunofluorescent staining was done to detect microglia. General microglia were identified by immunostaining for Ionized calcium-binding adaptor molecule 1 (Iba1) (chicken mAb anti-Iba1; #IBA1-0100, Aves Labs). The resident glia population was identified using TMEM119 Type (Guinea pig mAb anti-TMEM119 #400 308, Synaptic Systems). We used a free-floating section method for immunostaining. First brain sections were blocked in 5% normal goat serum in 0.1M phosphate buffer saline (PBS) supplemented with 0.3% Triton-X100 for one hour, then incubated overnight in a cold room on a shaker, in primary antibodies in PBS with 3% normal goat serum and 0.3% Triton-X100. Following that, sections were washed in 0.1M PBS, then incubated in goat anti-chicken IgY Alexa Fluor 488 and goat anti-guinea pig IgG Alexa Fluor 555. Lastly, the sections were rinsed in 0.1M PBS and incubated in DAPI (#62248, ThermoFisher Scientific), rinsed, and mounted on Superfrost Plus slides (#12–550-15, Fisher Scientific) in Prolong Diamond with DAPI (#P36966, ThermoFisher Scientific).

### Quantitative confocal microscopy

We quantified immunofluorescent cells in the cornu ammonis 1 (CA1) area of the hippocampus as described previously.^[75]^ We obtained confocal images (Leica TCS SP8 confocal microscope) at 1 μm steps through the z-axis, with Leica 63X×/0.75 (zoom factor of 0.75) objective. Counts of labeled microglia were made in 4 sections, spaced 400 μm apart, along the dorso-ventral axis of the hippocampus. Within the region of interest, counts were performed in optical sections spaced 1-μm apart (typically 18–20 optical sections along the z-plane.

### Immunoblotting

Sodium dodecyl sulfate-polyacrylamide gel electrophoresis (SDS-PAGE) was carried out as previously described.^[76, 77]^ Briefly, rapidly dissected cortical/hippocampal tissue from female WT and MeCP2^+/-^ mice was dissected and collected in a RIPA buffer. Samples were diluted in 2x sample buffer, and 30 μg of protein was loaded onto a 7% polyacrylamide gel. Next, proteins were transferred onto a nitrocellulose membrane, blocked, and probed with primary antibodies. The membranes were washed and incubated for 1 hour at room temperature with HRP-conjugated secondary antibodies. Protein bands were visualized with Pierce ECL and imaged using a ChemiDoc MP (Bio-Rad). Band intensity was compared to beta-actin and beta-tubulin as loading controls.

### Densitometry

For Western blot analysis, bands from raw images were analyzed using densitometry. Biological replicates were run on the same gels for comparison, and each band’s area under the curve was calculated. The average signal for each treatment group was calculated based on the protein expression levels.

### Statistical Analysis

All data are presented as the mean ± standard error of the mean (SEM). The Shapiro-Wilk test was performed to find the normal distribution of the data sets. EEG data were analyzed using the Welch’s t or Mann-Whitney t test. Bioavailability and open field data were analyzed using the Welch’s t-test. Sociability data was analyzed using the Welch’s t-test, and preference of sociability data was analyzed using the paired t-test. To analyze the novel object recognition task, we used Welch’s t-test and paired t-test. The hindlimb clasping data were analyzed using the Mann-Whitney t-test. The Neuroinflammation data were analyzed using the Welch’s t or Mann-Whitney t test. Two-way ANOVA followed by Tukey’s Multiple Comparison was used to test the western blotting data. P values < 0.05 are considered statistically significant.

All animals and reagents used for these studies are listed in Table S1: Key Resources.

## Author contributions

M.N.A., T.N., Z.Z., J.L.S., P.A.D., and S.J.M. designed research; M.N.A., S.F.J.N., S.S., and K.A, performed research; M.N.A., S.F.J.N., S.S., and K.A, analyzed data; M.N.A. and S.J.M. wrote the paper with input from all the authors.

## Data Availability

Data will be made available on request.

